# RdRp-scan: A Bioinformatic Resource to Identify and Annotate Divergent RNA Viruses in Metagenomic Sequence Data

**DOI:** 10.1101/2022.02.28.482397

**Authors:** Justine Charon, Jan P. Buchmann, Sabrina Sadiq, Edward C. Holmes

## Abstract

Despite a rapid expansion in the number of known RNA viruses following the advent of metagenomic sequencing, the identification and annotation of highly divergent RNA viruses remains challenging, particularly from poorly characterized hosts and environmental samples. Protein structures are more conserved than primary sequence data, such that structure-based comparisons provide an opportunity to reveal the viral “dusk matter”: viral sequences with low, but detectable, levels of sequence identity to known viruses with available protein structures. Here, we present a new open computational and resource – RdRp-scan – that contains a standardized bioinformatic toolkit to identify and annotate divergent RNA viruses in metagenomic sequence data based on the detection of RNA dependent RNA polymerase (RdRp) sequences. By combining RdRp-specific Hidden Markov models (HMM) and structural comparisons we show that RdRp-scan can efficiently detect RdRp sequences with identity levels as low as 10% to those from known viruses and not identifiable using standard sequence-to-sequence comparisons. In addition, to facilitate the annotation and placement of newly detected and divergent virus-like sequences into the known diversity of RNA viruses, RdRp-scan provides new custom and curated databases of viral RdRp sequences and core motif, as well as pre-built RdRp alignments. In parallel, our analysis of the sequence diversity detected by RdRp-scan revealed that while most of the taxonomically unassigned RdRps fell into pre-established clusters, some sequences cluster into potential new orders of RNA viruses related to the *Wolframvirales* and *Tolivirales*. Finally, a survey of the conserved A, B and C RdRp motifs within the RdRp-scan sequence database revealed additional variations of both sequence and position, which might provide new insights into the structure, function and evolution of viral RdRps.

## INTRODUCTION

The explosion of viral metagenomic projects and associated high-throughput sequencing data over the last decade have paved the way for a reappraisal of the diversity and evolution of RNA viruses^1–3^. The expansion of the RNA virus discovery to overlooked habitats, environments and organisms has led to the recognition that RNA viruses are ubiquitous and likely infect all types of cellular organisms^1,2,4–8^. Although there is an understandable focus on potential health impact of RNA viruses, their importance goes far beyond their role as pathogenic agents in humans, domestic animals and livestock. For example, investigating RNA virus diversity in marine habitats has revealed their importance in fundamental ecological and biogeochemical processes^9,10^. Attempts to illuminate the RNA virus world are also expected to provide fundamental insights into the origins of RNA viruses and the long-standing evolutionary processes that have shaped their current diversity.

RNA viruses exhibit the highest evolutionary rates of any organism^11,12^ and do not possess universally conserved and easily interpretable gene sequences equivalent to the 16S and 18S rRNA used to classify other microorganisms. The identification of RNA viruses from metagenomic data sets largely relies on sequence similarity-based comparisons. While such studies have enriched the RNA virus phylogeny, they are only able to detect sequences with relatively high levels of sequence similarity to existing sequences, with the detection usually limited to proteins sharing at least 30% sequence identity^13^. As a consequence, it is highly likely that a vast world of undiscovered RNA virus sequences - the so-called “viral dark matter”^3,14^ - is present within the sequence data generated to date but which are too divergent to detect with currently available bioinformatic tools.

Protein structures are up to ten times more conserved than nucleotide sequences^15^. Integrating protein structural-comparison steps into metagenomic pipelines is therefore expected to greatly extend our limit of detection^16,17^, potentially enabling the identification of sequences in the range of 10 to 30% identity (the protein “twilight zone”^13^) and for which corresponding viruses might be referred to as the “viral dusk matter”. The viral RNA-dependent RNA polymerase (RdRp or replicase) is the most conserved protein in RNA viruses^18–20^. As a consequence, the RdRp is widely used as a reference gene in the estimation of RNA virus phylogenies, which in turn form the basis of the newly-established classification scheme for RNA viruses^21^. Because of its central role in genome replication and transcription, the RdRp is essential from earliest stages of the virus infection cycle, and RdRps are relatively well characterized at both the functional and structural levels^18–20^.

Viral RdRp structures can be envisaged as comparing a “right hand” shape with thumb, palm and finger subdomains^19,22–24^. The thumb region helps in complex interactions and stabilization between the RNA template and free nucleotides, while the finger subdomain is both involved in recognition and binding to nucleic acids and positioning the template for polymerisation catalysis. The palm domain is the most conserved and forms the catalytic core of RdRp. Crucial residues and secondary structures are involved in this catalytic function, and three main amino acid motifs – denoted A, B and C – have been reported as consistently conserved and identified at both the primary and secondary structure levels.

The catalytic A motif is formed by an invariant aspartate as well as a second (Dx2-4D) in most of RdRps reported to date^20^. The second aspartate can be replaced by a Lysine residue in single-strand negative-sense (ss-) RNA viruses^20^. The B motif is involved in template binding and displays a conserved glycine to potentially allow conformational changes required to accommodate template and substrate interaction, and its sequence varies between the major groups of RNA viruses. The C motif is the most conserved. It comprises a loop that most commonly contains a triplet GDD motif flanked by two beta strands. Along with the aspartate of the A motif, it forms the catalytic triad essential for metal ion binding and coordinating the elongation reaction of the newly synthetized strand. Despite its strong conservation, sequence variation has been reported in some viral families with, for example, a GDD/ADN C motif preceding the A motif in the *Permutotetraviridae* and *Birnaviridae*, respectively^19,25–27^. Similarly, in some other single-strand positive-sense (ss+) and ss(-)RNA viruses, the glycine in the GDD motif has been replaced by a serine (S)^28,29^, while the second aspartate has been replaced by an asparagine (N) in some ss(-)RNA viruses^29,30^.

Recent studies have emphasized the conserved nature of RdRp protein structures and associated mechanisms of catalysis among RNA viruses as a whole, reinforcing the hypothesis of their ancient common ancestry, despite the very low levels of amino acid sequence similarity^18–20,31^. The structural conservation of the RNA virus RdRp and the resulting “evolutionary fingerprints” at the sequence levels therefore provide a valuable tool to infer remote protein homology and relatedness, such that the RdRp appears as the candidate of choice for the structure-based exploration of the viral dusk matter.

The use of Hidden Markov Models (HMMs), rather than primary sequences, also provides a powerful way to identify highly divergent viral sequences. By converting protein multiple sequence alignments (MSAs) into statistical models with position-specific scores, HMMs integrate the evolutionary information and have been shown as particularly powerful in the detection of remote protein homologies^32^. Importantly, HMMs tend to outperform classical sequence-based profiles or sequence alignment approaches, providing a more sensitive performance at minimum computational cost^32,33^ and therefore constitute a promising approach to improve the detection of divergent RNA viruses^16^.

Herein, we outline a new set of computational methodologies and resources to identify and annotate remote viral RdRp signals from metagenomic data sets. To do so, we first inferred the existing diversity of viral RdRp contained in the non-redundant (nr) database from the National Centre for Biotechnology Information (NCBI), constituting the largest protein database currently available (NCBI; https://www.ncbi.nlm.nih.gov/) as well as in some of the most recent RNA virus metagenomic surveys. By conducting filtering and curation steps, we developed a global and non-redundant RdRp core database that was then used for the description of the RdRp diversity and functional annotation among the already established clades of RNA viruses. The performance of some current tools to identify remote RdRp signals from primary sequence was then evaluated, and a new custom RdRp-profile database was created to significantly improve the profile-based homology detection. Finally, an overall profile and structural-based workflow was proposed.

## 2. MATERIALS AND METHODS

### 2.1 RdRp sequence database

#### Retrieval of viral RdRps using keywords

The *Riboviria* annotated RdRps contained in the NCBI non-redundant protein (nr) database (*https://www.ncbi.nlm.nih.gov/protein*) were searched using the Entrez Programming Utilities from the NCB*I* (*https://www.ncbi.nlm.nih.gov/books/NBK25501/*) and retained based on a 300 amino acid length cut-off and a list of RdRp-related keywords available at https://github.com/JustineCharon/RdRp-scan/rdrp_keyword.list. Proteins containing no-RdRp description were filtered out based on keywords (“polymerase cofactor”; “glycoprotein”; “capsid protein”; “subunit”; “nucleocapsid”; “NS5a”; “matrix”; “gglutinin”; “coat”; “reverse”). A 90% identity clustering was performed using CD-HIT (CD-HIT v4.6.1)^34^. Manual curation was conducted based on the presence of the tree canonical A, B and C motifs using the Geneious software (v11.1.4)^35^.

#### Addition of RdRp sequences retrieved from metagenomic studies

Additional metagenomic-based RdRp sequences were retrieved from recent studies^36,37^ and added to the nr RdRp using CD-hit 90% (v4.6.1)^34^.

#### Blast analysis of the nr database

*A* comparison against the nr NCBI database was performed using Diamond BLASTp (v2.0.9) ^38^(e-value cut-off 1e-05; “very-sensitive” option).

#### Taxonomic assignment of RdRps

NCBI taxonomy assignments were retrieved using the option lineage from the taxonkit tool available at https://github.com/shenwei356/taxonkit^39^. All NCBI RdRp sequences were grouped according to their placement within the realm *Riboviria*, kingdom *Orthornavirae* (which includes all the RdRp-encoding RNA viruses), using both order and phylum ranks according to the current ICTV virus classification (https://talk.ictvonline.org/taxonomy/). The clustering and automatic assignment was performed using cd-hit-2d with 60%, 50% and 40% successive clustering (v4.6.1)^34^. The 30% clustering of unclassified sequences was performed using hierarchical clustering, with the h3-cd-hit option of CD-HIT using the WebMGA server available at http://weizhong-lab.ucsd.edu/webMGA/server/psi-cd-hit-protein/. Briefly, h3-cd-hit performs three iterative runs of CD-HIT clustering at 90%, 60% and 30% identity, using a neighbour-joining method.

#### Host assignment

Whenever available, the putative host information of each RdRp-corresponding virus was retrieved using taxonkit name2taxid available at https://github.com/shenwei356/taxonkit39.

### 2.2 RdRp profile database

#### Palmdb RdRp core sequences

Palmdb RdRp sequences (uniques.fa - version 02/03/2021) and those generated by the Serratus project update (serratus.fa - version 14/03/2021) were both retrieved from the Palmdb github repository (https://github.com/rcedgar/palmdb).

#### RdRp multiple sequence alignment and profile construction

For each viral order and phylum as well as unassigned clusters, RdRp sequences were enriched with the Serratus/Palmdb RdRp sequences sharing more than 40% sequence identity using CD-HIT-2d (v4.6.1)^34^. Redundancy was removed at the 40% identity level using CD-HIT, and multiple sequence alignments (MSA) of various sizes were obtained using Clustal Omega (--auto option) (v1.2.4)^40^ (Figure S2). The resulting sequence alignments were manually curated to remove partial RdRp core sequences using Geneious software (v11.1.4)^35^. HMM-profiles were built from each MSA using HMMer3 using standard parameters (v3.3)^41^. The resulting HMM-profiles were combined and converted into one final HMM-profile database used in the subsequent profile analysis with the HMMer3 hmmpress option.

#### Profile construction of the unassigned viral RdRp

Clusters of “unassigned” sequences (i.e. sharing <30% sequence identity with RdRp members of pre-established viral groups) containing more than 10 sequences were enriched with the Serratus/Palmdb RdRp sequences sharing more than 40% sequence identity using CD-HIT-2d (v4.6.1)^34^.

### 2.3 Constructing a RdRp A, B and C motif database and phylogenetic analysis

Alignments of the RdRp were built for each virus phylum using Clustal Omega (--auto option) (v1.2.4)^40^ and the previously 40% sequence identity clustered sequence files, depleted of the Serratus and Palmdb sequences. RdRp A, B and C motif sequences were extracted from RdRp alignments and corresponding logos were obtained using WebLogo (v3.7.8)^42^. For phylogenetic analysis, iterative alignments of RdRp were processed using the -p1 option using a previous structural alignment^18^ as a backbone. Unclassified sequences were then aligned to the intermediary alignment, resulting in a final alignment of ~3,300 sequences (Figure S2). Intermediary and final alignments were manually inspected using Geneious software to check for the presence of aligned motif blocks. Finally, phylogenetic trees were inferred from this alignment using FastTree2 (v2.1.9)^43^, an approximate maximum likelihood-based method, applying the default options. Resulting phylogenies were mid-point rooted and represented using FigTree software.

### 2.4 Interproscan analysis

Our custom RdRp database was compared to the EBI InterPro-embedded PANTHER, Pfam, PIRSF, PRINTS, PROSITEPATTERNS, PROSITEPROFILES, SUPERFAMILY and TIGRFAM protein profile databases using Interproscan (v5.52-86.0)^44^. RdRp-like profile entries are listed in (https://github.com/JustineCharon/RdRp-scan/RdRp_InterPro_keyword.list).

### 2.5 Phyre2 analysis

Analyses of structural homology were conducted using the batch mode of the Phyre2 server^45^, available at http://www.sbg.bio.ic.ac.uk/phyre2/html/page.cgi?id=index. The pre-clustering of RdRp sequences at 30% identity were performed using the h3-cd-hit option of the WebMGA server available at http://weizhong-lab.ucsd.edu/webMGA/server/psi-cd-hit-protein/.

### 2.6 RdRp-scan workflow

Open reading frames of orphan contigs were obtained using the GetORF tool from the EMBOSS package (v6.6.0)^46^ using the -find 0 option (defining an ORF as sequence between two STOP codons). The list of genetic codes used by viruses was retrieved from the NCBI Taxonomy resource (https://www.ncbi.nlm.nih.gov/Taxonomy/CommonTree/wwwcmt.cgi) and corresponding translation tables 1,3,4,5,6,11 and 16 used to translate potential viral ORFs.

## 3. RESULTS and DISCUSSION

### 3.1 A new curated viral RdRp database

Revealing the diversity of RNA viruses in metagenomic data sets relies on our ability to compare sequenced and assembled contigs to pre-existing viral nucleotide and protein databases. However, current general protein databases are either too large (NCBI nr) or not comprehensive (NCBI RefSeq), while viral-specific ones such as the Reference Viral Database (RVDB)^47,48^ also contain DNA viruses and endogenous virus sequences. In addition, many RdRp-like sequences obtained from viral metagenomic studies are mis-annotated or not assigned to any function and/or viral clades. Such factors can compromise the detection of more divergent viral sequences and slow analyses by requiring extensive computational resources or providing high numbers of false positive-results (*e.g*. non-RNA virus hits). To facilitate and speed the specific detection of RNA virus signals, we provide a comprehensive non-redundant and manually curated viral RdRp protein sequence database. Specifically, we compiled into one location all the RdRp and RdRp-like sequences contained in the nr NCBI protein database, as well as from two major viral metatranscriptomic studies that identified thousands of new RNA viruses^2,37^.

#### A new viral RdRp database

To build our viral RdRp-specific database, we first manually retrieved all the RNA virus encoded RdRps from the nr NCBI database (*https://www.ncbi.nlm.nih.gov/protein*) based on taxonomy (Realm: Riboviria - Kingdom: *Orthornavirae* - excluding *Pararnavirae* retro-viruses), keywords and a length cut-off of 300 amino acids (Figure 1). The RdRp sequences from recent metagenomic studies^2,37^ were added, and a filtering and manual curation step was employed to remove all non-RdRp sequences. Briefly, the “core” region of the RdRp (i.e., containing the three motifs A, B and C) was located and extracted based on the identification of the A, B and C sequence motifs. RdRps without identifiable C motifs or displaying non-RdRp domains were removed. Sequences in which A and B motifs could not be directly identified were functionally annotated using the EBI Interproscan package^44^ to check for the presence and the position of an RdRp-like sequence. To retrieve additional unannotated RdRp and facilitate further alignment and annotation steps, only the RdRp core region and partial N-terminal and C-terminal flanking regions were retained, using an arbitrary length cut-off of 600 amino acids. Finally, sequences with>90% sequence identity to each other or known sequences were excluded to facilitate manual curation (Figure 1).

**Figure 1.**
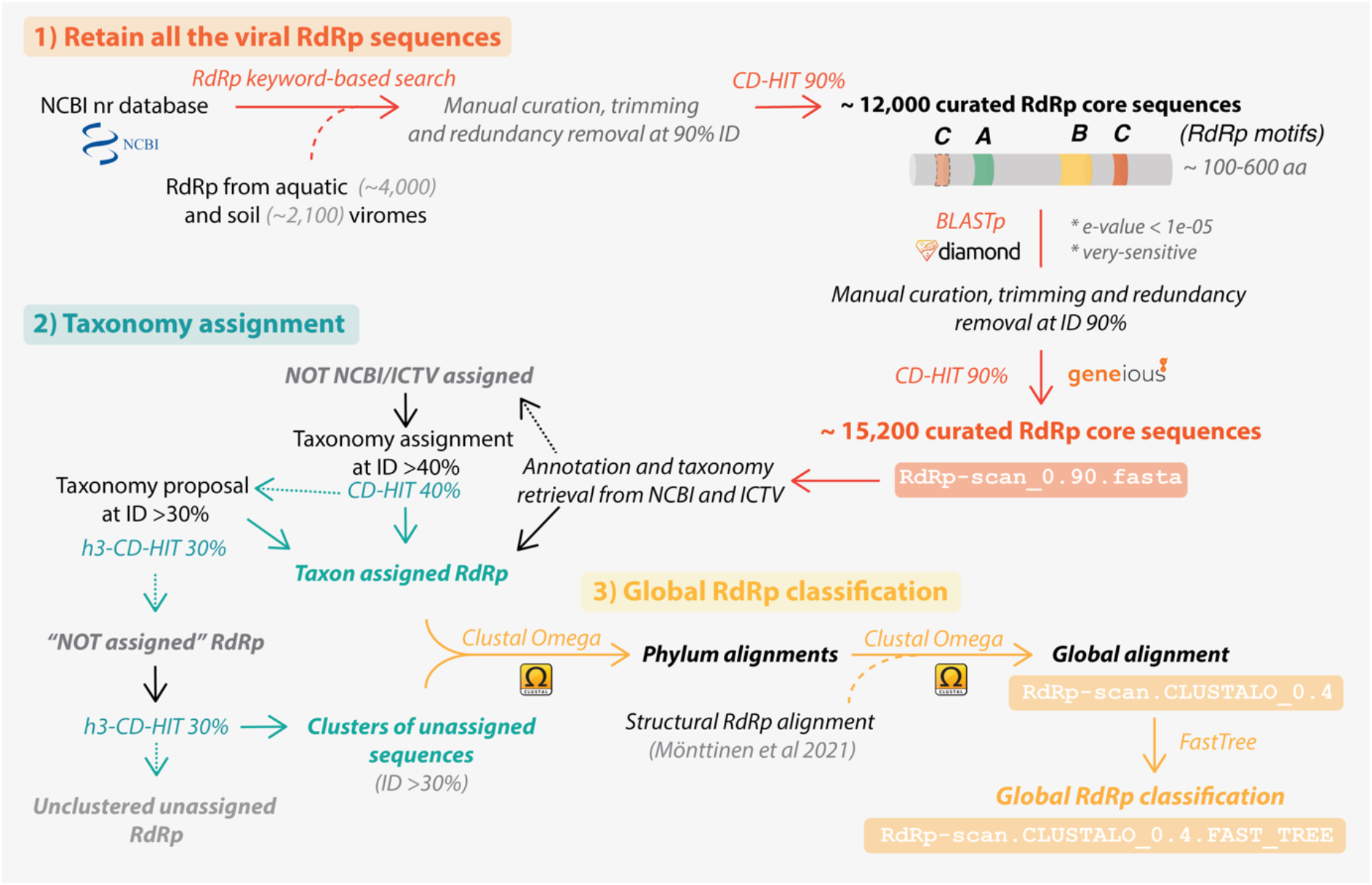
RdRp-scan sequence database workflow. Description of the procedure used to build the RdRp-scan RdRp sequence database and associated annotations. The file names are indicated in orange and yellow boxes and available at https://github.com/JustineCharon/RdRp-scan.

A BLAST search against the nr database was then conducted using the curated RdRp core sequences to ensure database completeness. All the RdRp-like homologs were retrieved, manually checked, trimmed and their redundancy removed as described previously. The NCBI information (taxonomy and host when available) of every nr-based sequence was retrieved and used to classify proteins based on the corresponding RNA virus phyla or orders (Figure 1). An attempt was made to automatically assign viral RdRp sequences to pre-established virus orders based on an arbitrary 40% sequence identity cut-off, widely used in protein sequence analysis^49^. A second clustering at 30% sequence identity was also used to suggest a taxonomic assignment at the order rank level for viruses that contain more divergent RdRps. Whenever possible, the final non-assigned RdRp core sequences (i.e., sharing <30% identity with any of the assigned RdRp) were clustered into “unassigned clusters” using a 30% identity cut-off (Figure 1). The resulting sequence database of RdRp clustered at 90% sequence identity is available at https://github.com/JustineCharon/RdRp-scan/.

To describe newly-identified RdRps, and potentially suggest a taxonomic placement, intermediary alignments of the *Riboviria* were built at both phylum and order levels. Considering the very high level of sequence divergence between the phyla of RNA viruses, we used a previous structural alignment^18^ as a backbone, and pre-aligned RdRps were iteratively added to build the final alignment (Figure 1). Finally, unassigned sequences were added to the master RdRp alignments. Importantly, at this level of divergence, the quality of the resulting alignments cannot be used to accurately infer RNA virus phylogenies. Rather, they should be regarded as broad indicators of how the unassigned sequences, and ultimately any newly discovered sequences, fit into global virus diversity.

#### Overall RNA virus diversity

The RdRp database provided in this study was also designed to infer the diversity of RdRp sequences covered by the current nr database. The distribution of all the curated RdRp sequences obtained at the 90% sequence identity level reveals that our global knowledge of RdRp diversity is clearly heterogeneous, partial and subject to profound sampling biases. This is particularly obvious at the level of virus hosts, with mammals and land plant-infecting virus RdRps accounting for almost 80% of all the host-assigned viruses (Figure 2). The very limited proportion of RdRp from RNA viruses infecting Archaea and microbial eukaryotes highlights the clear need to investigate viral diversity in those hosts (Figure 2). The bias in the distribution of RdRp diversity is also apparent in phylogenetic analyses in which those viral clades associated with mammals (e.g,. *Picornavirales)* and land plants (e.g., *Tolivirales)* are over-represented (Figure 3).

**Figure 2.**
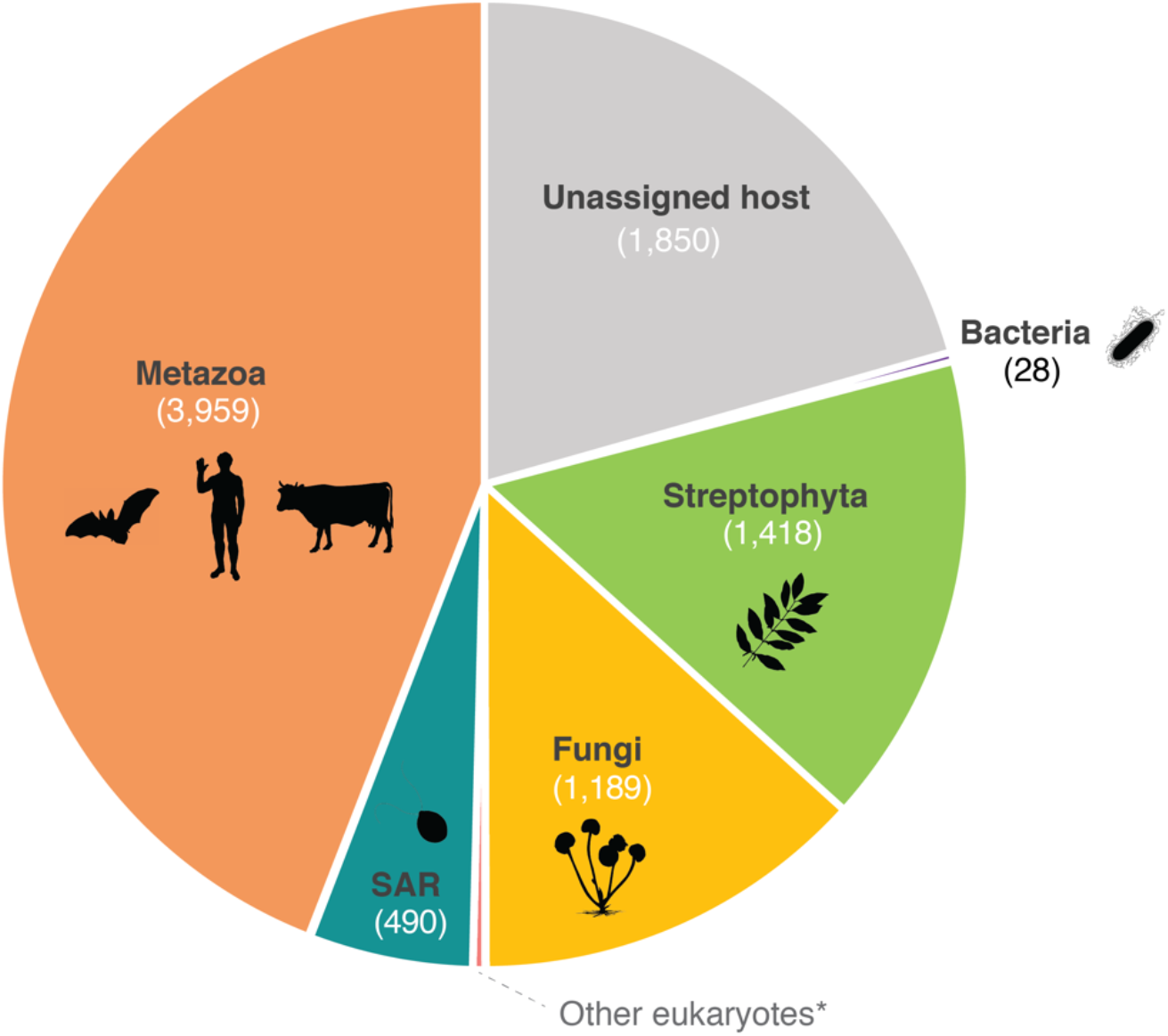
Host assignment of RdRp corresponding viruses. Host information was retrieved for each RdRp viral entry at NCBI present in the 90% redundancy RdRp-scan database, using either the VirusHostdb ^50^ and NCBI resources (https://www.ncbi.nlm.nih.gov/). Archaea were not represented as no archaea-infecting RNA viruses have been formally reported. *Other eukaryotic clades: Picozoa, Chlorophyta, Rhodophyta, Metamonada, Kinetoplastea, Amoebozoa, Haptophyta. Icons were retrieved from Phylopic (http://phylopic.org/). Credits: Matt Crook (Bacteria), Sergio A. Muñoz-Gómez (SAR), Ville Koistinen and T. Michael Keesey (Streptophyta), under the Creative Commons Attribution-ShareAlike 3.0 Unported license (https://creativecommons.org/licenses/by-sa/3.0/).

**Figure 3.**
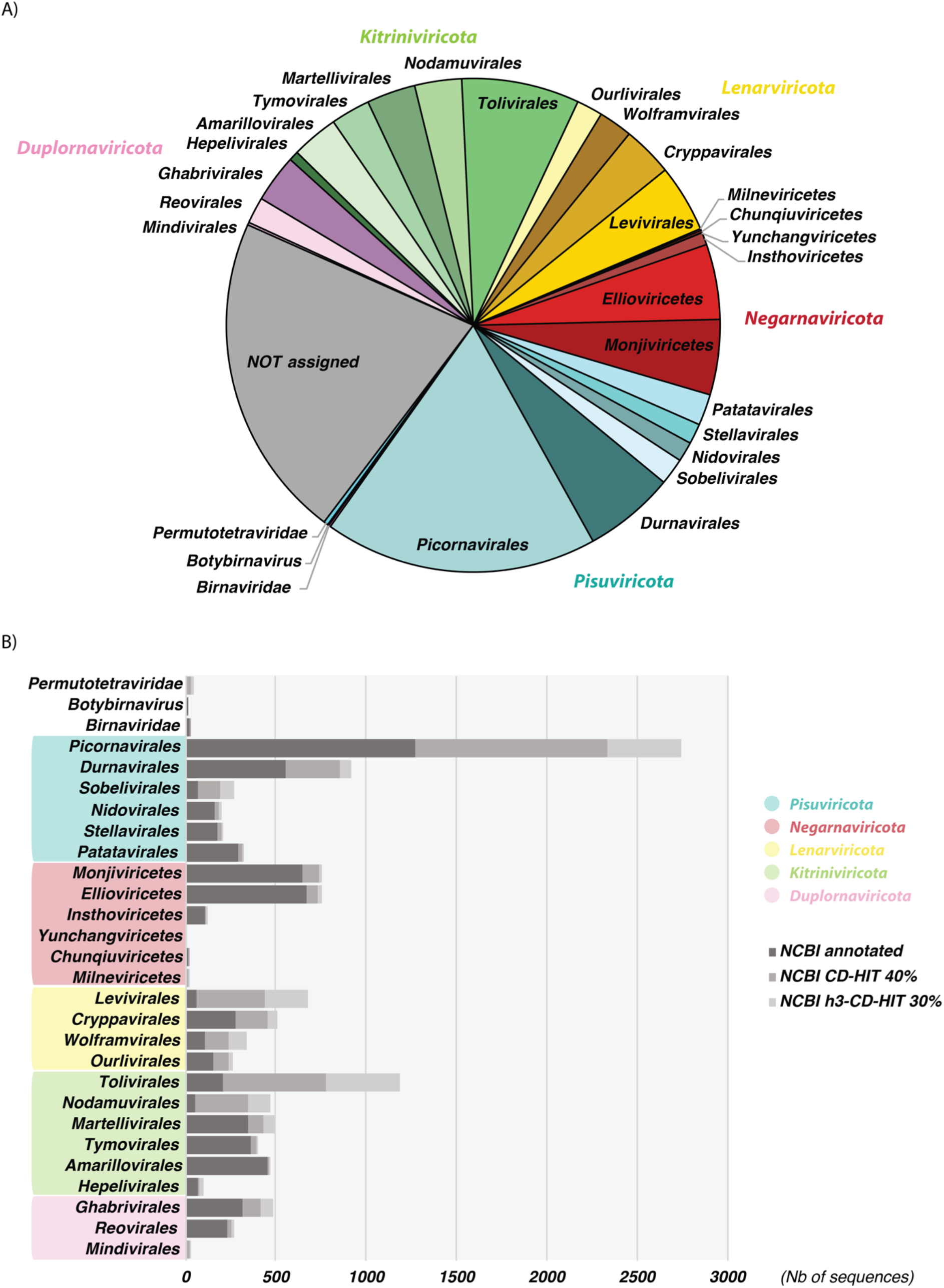
Taxonomic distribution of RdRps among the Riboviria phylum and clades. **(A)** Distribution of taxonomically assigned RdRp sequences (at 90% redundancy) among the taxonomy of the *Riboviria* versus those RdRp that are currently unassigned. **(B)** Distribution of NCBI annotated RdRp (dark grey) and automatically assigned RdRps at the 40% and 30% sequence identity levels (grey and light grey, respectively) among the current ICTV taxonomy of the *Riboviria*.

Interestingly, the phylum Lenarviricota encompassing viruses identified as infecting bacteria, fungi and unicellular eukaryotes (“protists”) from such families as the *Leviviridae, Mitoviridae* and *Narnaviridae*, contains a high number of sequences, most automatically assigned based on 30 to 40% sequence identity. Indeed, our automatic classification of previously unassigned sequences (i.e., showing >30% identity with assigned taxa) shows that phyla like the *Lenarviricota* are particularly enriched with such “assignable” sequences, and that their real diversity/importance may have been substantially underestimated.

We next attempted to integrate the unassigned sequences (i.e., those with <30% sequence identity with assigned RdRps) into the global RdRp phylogenies. This revealed that a large majority of the unassigned RdRps fell into pre-established families or clades with a reasonable degree of certainty (Figure 4). Moreover, the sequences that remain unassigned are distributed throughout global RdRp diversity, and their assignment may considerably enrich the diversity inside each RNA virus phylum. Only 24 RdRp were positioned outside of the currently established *Riboviria* phyla and without close assigned relatives. Such a small number is presumably a consequence of the biased sampling of viruses and the use of sequence-to-sequence tools in the metagenomic studies (Figure 4). Nevertheless, some major unassigned clusters largely obtained from metagenomic studies conducted on little studied environments (marine, soils) or hosts (unicellular eukaryotes, fungi), can be identified as distant relatives to the orders *Wolframvirales* (clusters 724, 295, 561, 197, 1028, 297 and 298) and *Tolivirales* (clusters 1279, 652 and more distantly clusters 1260, 731, 1282, 955, 589, 1135, 441, 373, 504 and 1039) and might constitute new orders within the *Lenarviricota* and *Kitrinoviricota* phyla, respectively (Figure 4). This strongly supports the notion that most of the RNA virus diversity remains uncharacterized, particularly at levels <30% sequence identity.

**Figure 4.**
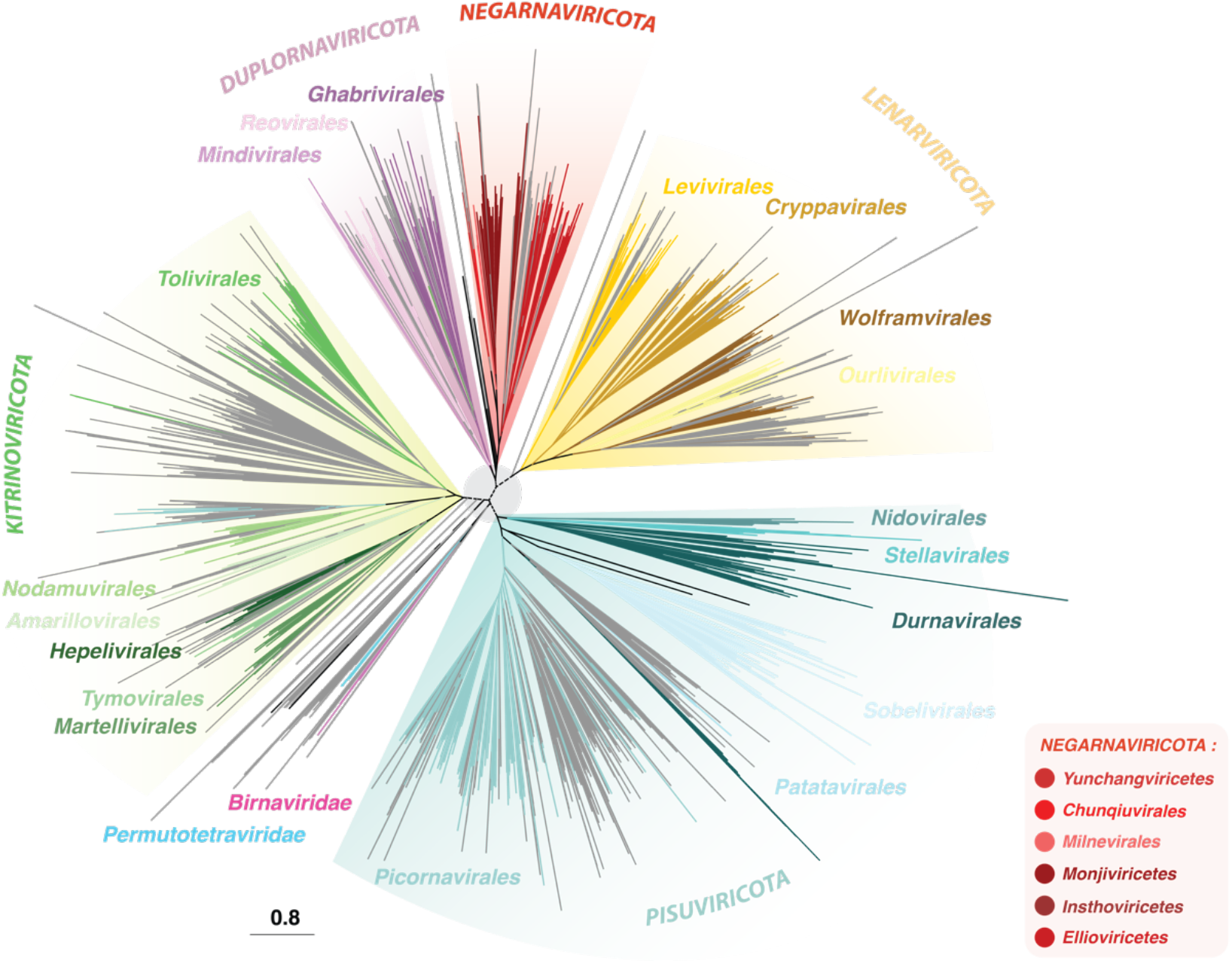
Unrooted phylogenetic tree obtained from the RdRp-scan protein sequence database. Given the high level of sequence divergence, the tree is unrooted, and a light grey circle is represented to highlight the uncertainty of the phylogeny at the inter-phylum level. *Riboviria* phyla *Pisuviricota, Kitrinoviricota, Duplornaviricota, Negarnaviricota* and *Lenarnaviricota* are coloured in blue, green, purple, red and yellow, respectively.

### 3.2 Identifying divergent viruses using profiles and structure-based homology

#### Ability of Interproscan and Phyre2 to detect divergent RdRps using our database

The capacity to detect RdRps using the currently available profile and structural databases was evaluated using two complementary approaches. First, Interproscan was run on our new custom RdRp database and the total proportion and number of detected RdRps was reported for each viral order (Figure 5). This revealed that Interproscan and integrated profile databases have a high capacity to detect RdRp sequences, with 97% of total RdRp detected regardless of viral taxonomy. Nevertheless, some viral orders were not well covered, with 18 to 85% of undetected RdRps in the *Wolframvirales, Ourlivirales, Reovirales* and *Chunqiuvirales* - or even completely missed in the case of the *Yunchangviricetes*.

**Figure 5.**
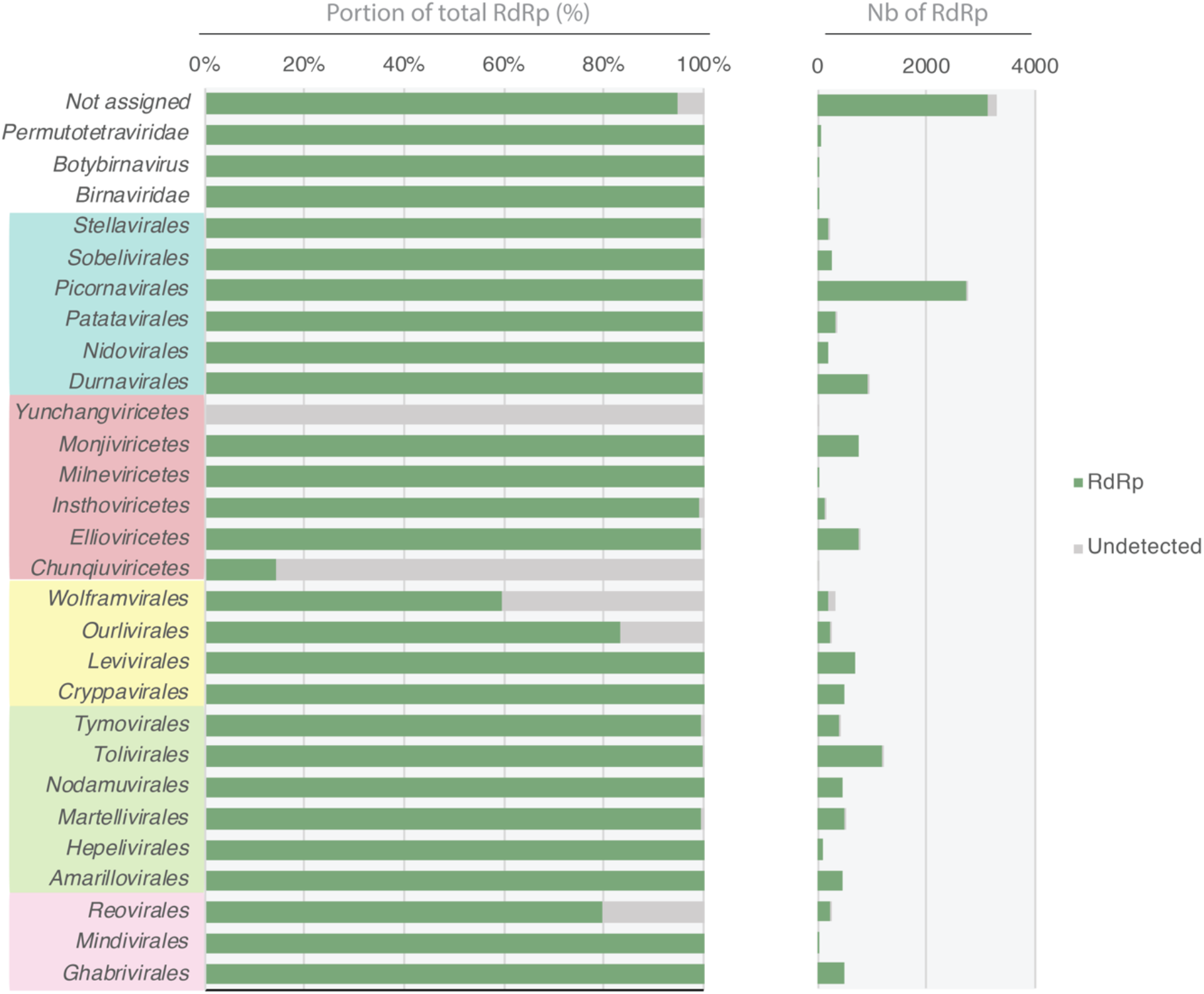
Detection of viral RdRp homology signal using Interproscan. The proportion (left part) and total numbers (right part) of RdRp detected and not detected by Interproscan are represented in green and grey, respectively. Viral orders are coloured according to their corresponding phyla and *Pisuviricota, Kitrinoviricota, Duplornaviricota, Negarnaviricota* and *Lenarnaviricota* are coloured in light blue, green, purple, red and yellow, respectively.

Next, to estimate the power of currently available tools to detect structural homology, we performed a comparison against the Protein Data Bank (PDB) (https://www.rcsb.org/)^51^ using the Phyre2 server^45^. Given the computational constraints inherent in Phyre2, a preliminary clustering of the RdRp at 30% identity was performed and only one representative sequence per cluster submitted to Phyre2, corresponding to around 1,500 sequences in total. We assume that protein sequences sharing more than 30% identity would have the same result at the level of protein structure comparison and that such clustering step does not affect the final results.

Remarkably, >96% of the RdRps submitted to Phyre2 were detected as homologous to the RdRp deposited in the PDB, regardless of viral phyla (Figure 6A). A limited number of RdRps could not be detected in few viral orders (*Wolframvirales, Ourlivirales, Durnavirales, Tolivirales* and *Reovirales*) or among the unassigned sequences (Figure 6A).

**Figure 6.**
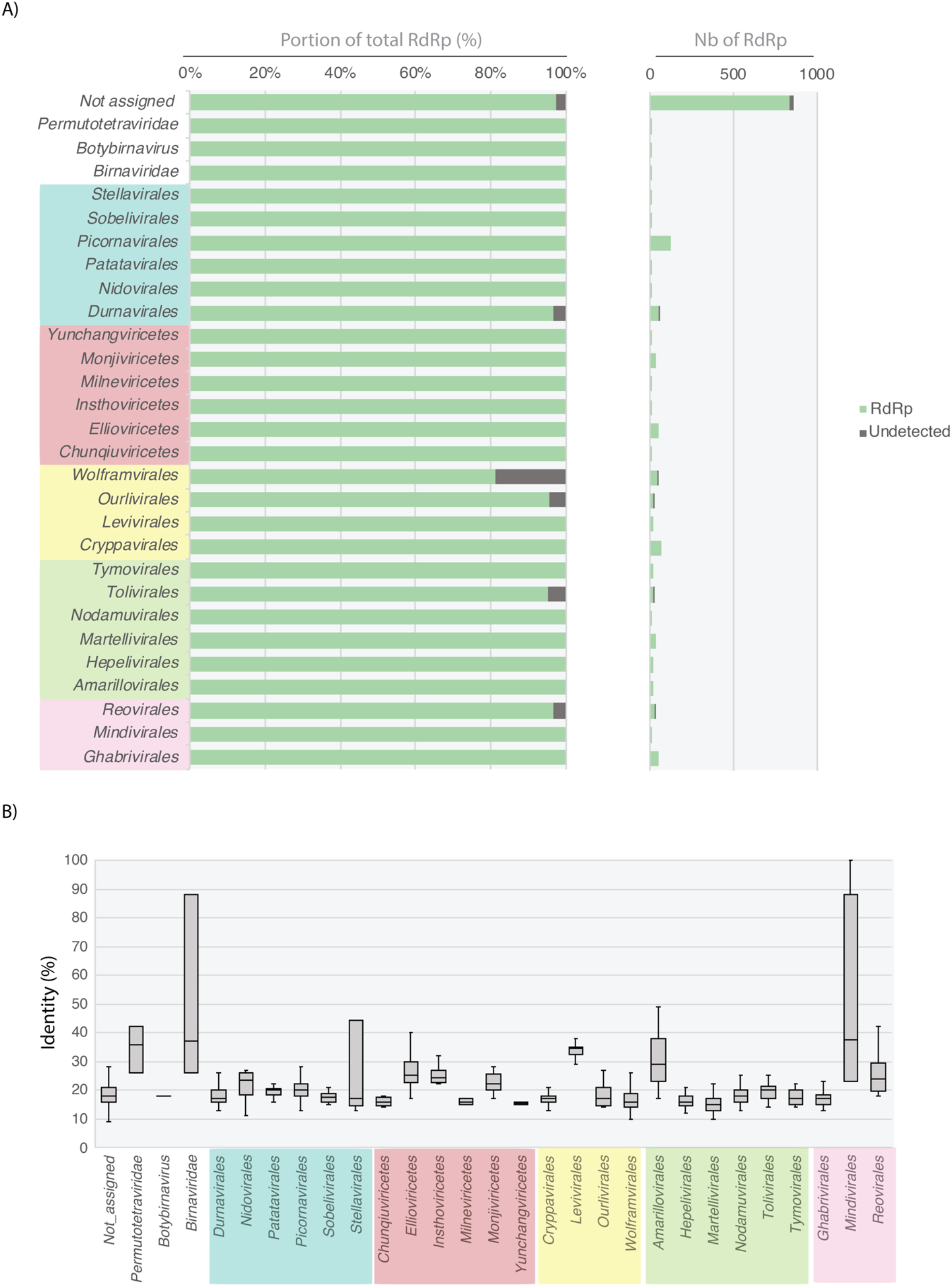
Detection of viral RdRp homology signal using Phyre2. (A) Proportion (left) and total numbers (right) of RdRps with PDB RdRp homolog detected at >80% of confidence levels for each order of the *Riboviria*. (B) Levels of sequence identity between Phyre2-detected homologs. The proportion (left) and total numbers (rightt) of RdRp detected and not detected by Phyre2 are represented in green and dark grey, respectively. Viral orders are coloured according to their corresponding phyla and *Pisuviricota, Kitrinoviricota, Duplornaviricota, Negarnaviricota* and *Lenarnaviricota* are coloured in light blue, green, purple, red and yellow, respectively.

Importantly, therefore, this analysis shows that Phyre2 can confidently detect homology at very low levels of sequence identity, with some of the PDB RdRp hits sharing as low as 10% identity with the RdRp sequences (Figure 6B). Despite the very limited representation of viral RdRps in the current PDB database^16^, Phyre2 is still capable of detecting RdRps by confidently identifying remote homologies between proteins that share very low levels of identity, with most <20%. As such, it constitutes a useful tool for the analysis of the viral dusk matter, although computational constraints make it difficult to use for most metagenomic data. In contrast, Interproscan processes thousands of RdRp sequences but fails to detect RdRp signals in some viral lineages. In addition, as it is not specifically designed for RNA viruses, the retrieval of all the different viral RdRp-specific signals needs to be performed manually using different keywords covering all the diverse viral RdRp-like profile names. Nevertheless, the use of protein HMMs is expected to help reveal deep evolutionary fingerprints shared between distant RdRp based on amino acid sequences.

#### Profile construction from the new RdRp db

To increase the diversity and specificity covered by RdRp-profile analyses and facilitate the detection of divergent viral RdRp homology signals, we built new RdRp profiles specific to each order or phyla of *Riboviria* based RdRp diversity obtained from both our RdRp database and those retrieved from unassigned RNA virus RdRp sequences. To maximize the sequence diversity covered by those new RdRp HMM-profiles, RdRp sequences recently obtained from the ultra-wide Serratus SRA mining project and members of the Palmdb resource^52,53^ were also integrated to pre-existing taxa using a 30% ID cut-off (Figure 7).

**Figure 7.**
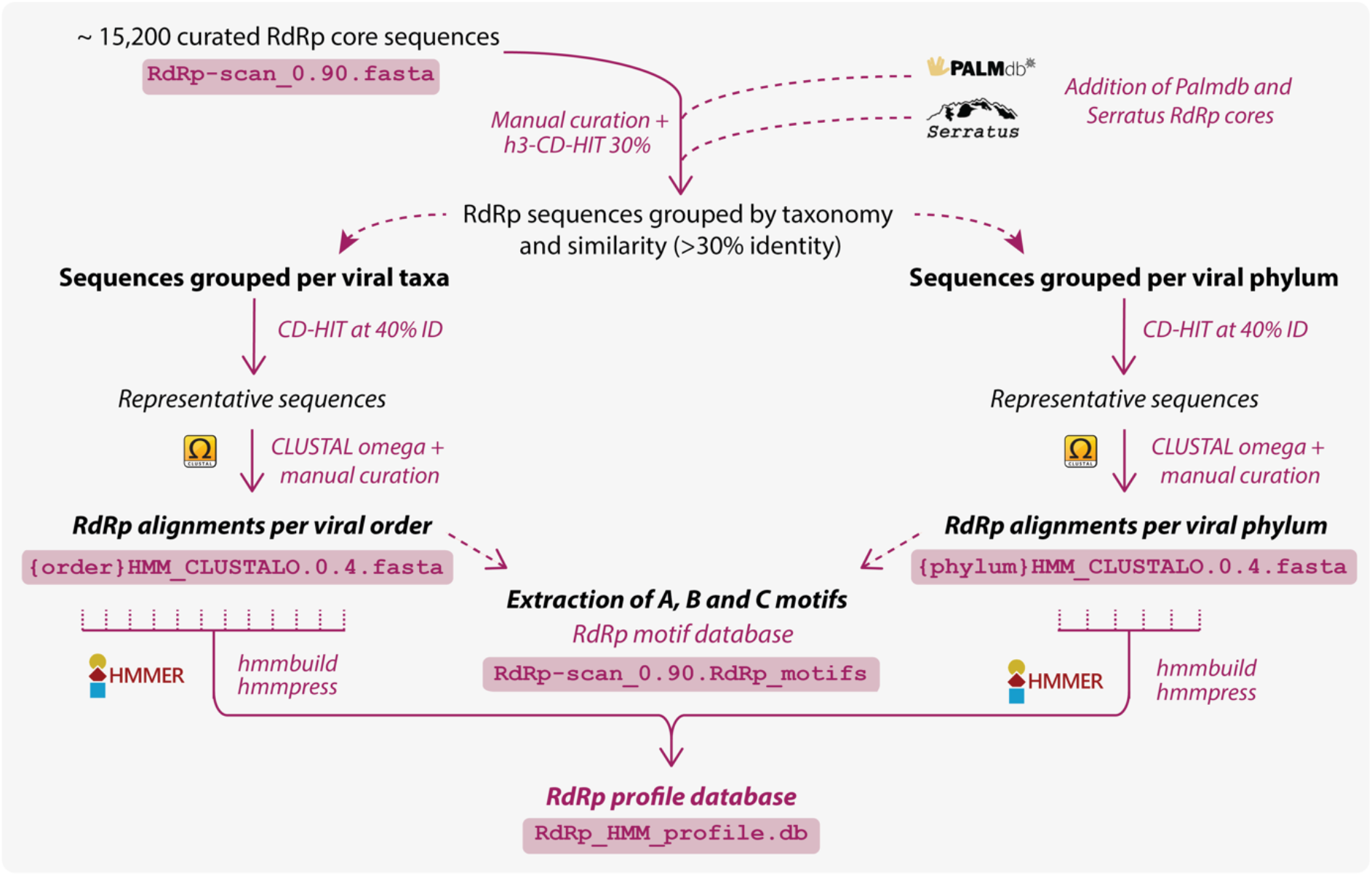
RdRp-scan profile database workflow. Description of the procedure used to obtain the HMM-based RdRp profiles. Corresponding file names are indicated in pink boxes and available at https://github.com/JustineCharon/RdRp-scan.

Each RdRp from our database as well as those from the Serratus SRA mining were grouped into viral orders or phyla according to their NCBI taxonomy assignment or their level of similarity with taxonomically assigned RdRps, as described previously. Sequence alignments were produced for each group as well as for unassigned sequences after removing the redundancy at 40% ID. Each alignment was manually checked for the presence of A, B and C motifs, and whenever identified, sequences containing C motifs in N-terminal position were extracted and analysed separately. In total, 68 HMM-profiles were obtained (8 and 29 from established phyla and orders, respectively, and 31 from unassigned clusters) and then pooled into one single HMM-profile database to use with the HMMer3 package available at https://github.com/JustineCharon/RdRp-scan (Figure 7).

#### Catalytic motif diversity from the RdRp profiles

Covering the diversity of RdRp motifs is crucial to validate newly identified divergent candidates based on structure and profile-based homology. The A, B and C motifs were therefore manually identified from alignments used to build HMM-profiles (Figure 7) and corresponding motif sequence conservation were visualized using logo representations (Figure 8 and S2). To prevent mis-annotation, motif characterisation and RdRp validation were conducted exclusively based on those reported and/or validated from previous studies. Therefore, the existence of additional A, B and C motifs in viral RdRps not present in our database cannot be excluded.

**Figure 8.**
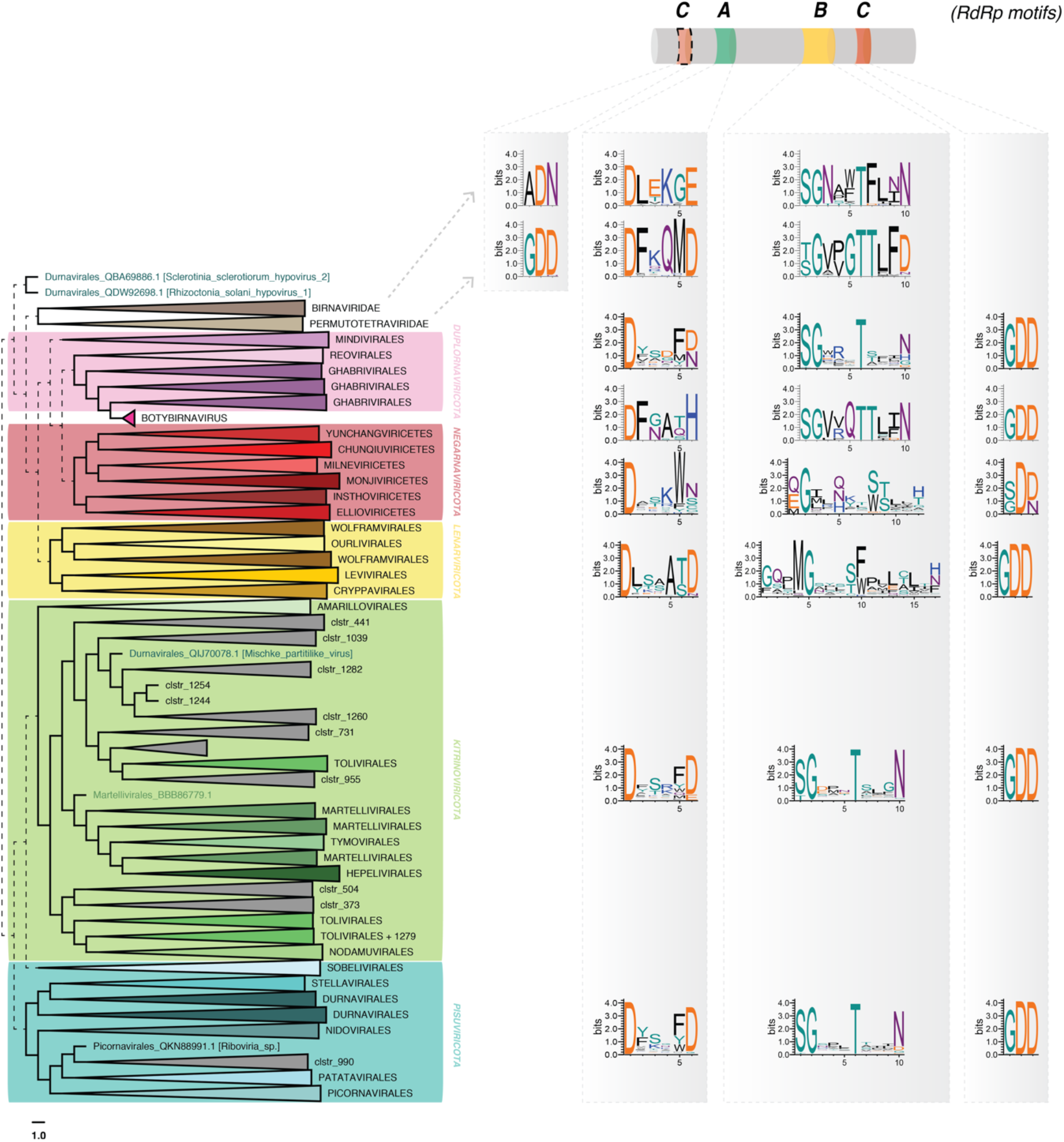
A, B and C RdRp motif diversity within *Riboviria* (*Orthornavirae*) phyla. Motifs are represented as logos obtained from alignment of RdRp within each phylum, using WebLogo ^42^. Dashed lines represent the uncertain placement/phylogenies due to the high level of divergence.

This analysis provided a number of notable observations. The A motifs show slight variation to the commonly described DxxxxD motif at both the phylum and order levels (Figures 8 and S2). While most of the *Pisuviricota* and *Kitrinoviricota* display a double aspartate residue, the second aspartate is replaced by E or H in the *Birnaviridae* and the genus *Botybirnavirus*. Members of the *Lenarviricota* commonly display an additional residue between the two aspartate (DxxxxxD), while the *Negarnaviricota* have a conserved tryptophan downstream of the critical aspartate residue (DxxxW) (Figures 8 and S2). The B motifs could be easily identified, and B motif sequences retrieved from the *Birnaviridae, Permutotetraviridae, Botybirnavirus, Duplornaviricota, Kitrinoviricota* and *Pisuviricota* based on the canonical SGxxxTxxxN, with only few variations at the level of virus order (Figure S2).

Conversely, the *Lenarvicota* display a very different and low-conserved motif at the primary sequence level, with a GQxMGxxxF/WxxL/ExxxF/H/N consensus. Similarly, the *Negarnaviricota* also display alternative B motif sequences, with a high diversity of residues arranged around the conserved glycine, crucial for the RdRp structure and function^20^. The organization and structural relevance of such alternative B motifs will need to be further investigated, but may provide insights into the structural and evolution of viral RdRps.

Of most note, variations were apparent in the C motif even though this is considered to be the most conserved RdRp motif. The GDD motif contains two of the three aspartate residues needed for RdRp catalytic functions^26^. As confirmed here, the C motif is most commonly observed downstream to the B motif, but exceptions have been reported with in *Birnavirus* and the permutotetraviruses that display a C-A-B motif order^25,54^. Moreover, the *Birnavirus* C motif comprises the ADN residue instead of the widely conserved GDD, replacing the third aspartate of the catalytic triad, which decreases the replicase activity^27^. Similar C-A-B organizations are newly reported here in the *Tolivirales, Martellivirales* (*Kitrinoviricota*) and *Patatavirales*, and could suggest a wider use of this uncommon C motif organization. While the GDN and SDD alternative sequences were already reported for C motif of the ss(-)RNA virus RdRps^29,30^, RdRps from the chunqiuviricetes show a conserved IDD. Importantly, however, these new variant consensus sequences or motif arrangements need to be experimentally validated and their role in RdRp structure and function investigated in more detail. Despite this, their identification will assist in the manual annotation of divergent viral RdRps and discriminate real exogenous viral signals from EVEs or other non-RdRp signals. Accordingly, a database of all the motifs identified in the RdRP db is provided at https://github.com/JustineCharon/RdRp-scan to help with such identification and the annotation of new viral candidates (Figure 7).

#### Proof-of-concept of RdRp detection using the MMETSP data set

To evaluate the performance of the new HMM-RdRp profiles in the detection of remote RdRp signals from meta-transcriptomic studies, unicellular eukaryotes transcriptomes from the Marine Microbial Eukaryote Transcriptome Sequencing Project (MMETSP)^55^ were used.

The MMETSP-based orphan ORFs (i.e., without detectable match in nr protein database using BLAST) obtained a previous study^5^ but not yet included in the current RdRp-scan database were submitted in parallel to HMMer3 using our HMM-profiles and to Interproscan using corresponding profile database. The resulting hits were then submitted to Phyre2 and a further validation using motif detection was performed. Sequences without motif similarities with known RdRp and/or detected as non-RdRp in Phyre2 with a confidence score higher than 90% were considered as false-positives. Previously identified divergent viruses that could not be detected using Interproscan or RdRp-scan profiles were reported as false-negatives. Previously identified viruses detected as true hits in this study, and true-like RdRp hits originally discovered using profiles were counted as true positives. Finally, the false-positive hits detected using one method but not detected using the other method were reported as “true negatives”.

Overall, the HMM-profile-based detection using our newly built RdRp profile displays a greater sensitivity with a significantly lower number of false-positives than Interproscan (Figure 9). Importantly, Interproscan was unable to detect five of the 30 previously reported RdRp sequences^5^. Conversely, by using HMMer3 combined to our new HMM-RdRp profile database, we were able to identify all the divergent viruses as well as new RdRp-like sequences. This result constitutes a proof-of-concept of the relevance of using a specific RdRp profile to detect remote viral signals. Among the ten RdRp-like hits identified only using HMM-profile database, five were similar to *Wolframvirales*, three were similar to *Ghabrivirales*, one matched with *Permutotetraviridae* and one was similar to *Chunqiuviricetes*. This is in accordance with the previously assessed Interproscan coverage of the RdRp diversity (Figure 5).

**Figure 9.**
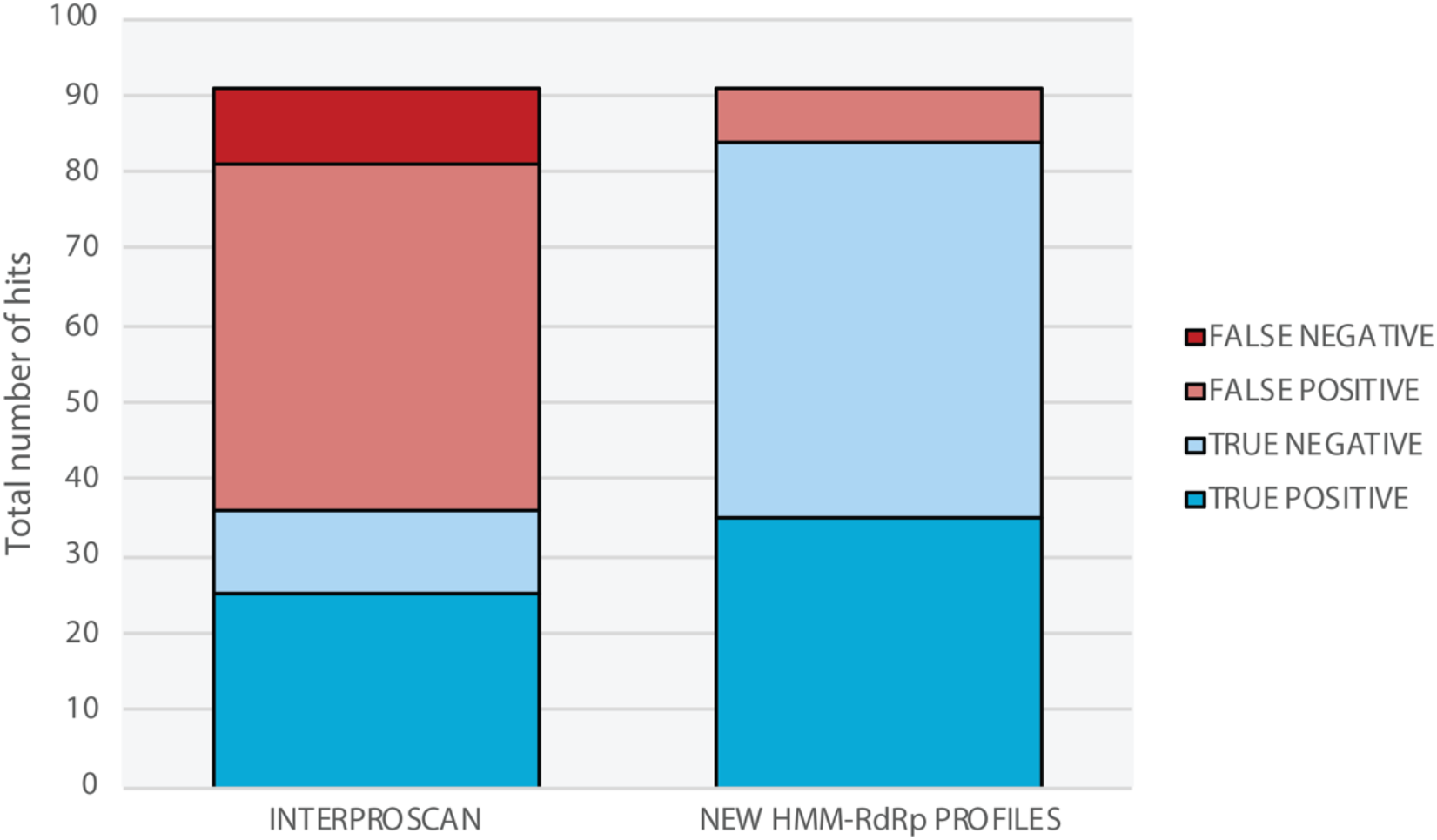
Comparison of RdRp detection from unicellular eukaryote transcriptomes using RdRp-scan HMM profiles versus Interproscan. Data sets obtained from MMETSP transcriptomes^5^ were submitted to HMMer3 using either RdRp-scan or Interproscan profiles. False-positives are indicated in light red and refer to sequences without RdRp A, B and C motif detected and/or detected as non-RdRp in Phyre2 with a confidence score higher than 90%. False-negatives are indicated in dark red and refer to previously detected viruses^5^ detected using Interproscan or RdRp-scan profiles. True positives are indicated in dark blue and refer to both previously detected viruses^5^ and true-like RdRp hits originally discovered using RdRp-scan and Interproscan profiles. True negatives are indicated in light blue and refer to false-positive hits detected using one method but not detected using the other method.

### 3.3 Analysis of new divergent RdRps using RdRp-scan

Together with the identification of divergent RdRp-like signals from metagenomic data, our study assists in the validation and characterization of newly identified RdRp-like sequences. Sequence comparisons with pre-existing RdRp can be challenging at such very low levels of similarity. Despite this, particular care should be applied when submitting new sequences, especially concerning: (i) the annotation of protein sequence(s) and (ii) the taxonomic placement of the corresponding virus within current viral diversity. While sequence annotation and taxonomic placement within the whole viral diversity is relatively straightforward for well-established families, it becomes a major challenge for new divergent sequences identified either from overlooked hosts or likely belonging to new/undescribed viral taxa that are prone to mis-classification.

#### Motif annotation using the motif database extracted from our custom database

To help in the annotation and detection of RdRp putative functional motifs from those divergent sequences, we have made available the RdRp motif databank extracted from the whole RdRp database which can be mapped directly onto candidate sequences using classical sequence visualization software (CLC Bio, Geneious, DNASTAR etc…). The three RdRp motifs A, B and C have a strong conservation and consistency among current RdRp diversity at both phylum and order levels (Figures 8 and S2). It is of obvious importance to verify the presence of the three motifs to prevent the mis-annotation of host genes as RdRps. Such RdRp-like signals identified in hosts always miss at least one of the three motifs, and this can be used to identify such false-positives. Profile alignments used to build the HMM-RdRp alignment are available at https://github.com/JustineCharon/RdRp-scan and can be used to align the candidate sequence to help identify conserved regions and motifs.

#### Taxonomic placement of new viral candidates

RdRp amino acid sequences are commonly used to infer RNA virus phylogenies. When attempting to taxonomically assign newly discovered sequences it is therefore tempting to put new virus candidates into the global diversity of RdRps. However, such large-scale RdRp phylogenies are not based on robust alignments. We therefore utilised the meta-RdRp alignment available at https://github.com/JustineCharon/RdRp-scan as an intermediary step to place the candidate sequence(s) within global RdRp diversity and identify their closest related sequences (Figure 10). The scale of analysis can then be narrowed, and the candidate sequences compared in a more detailed manner to the closest homologs (Figure 10). Reasonably robust phylogenetic trees and taxonomic placements can be obtained from these defined data subsets.

**Figure 10.**
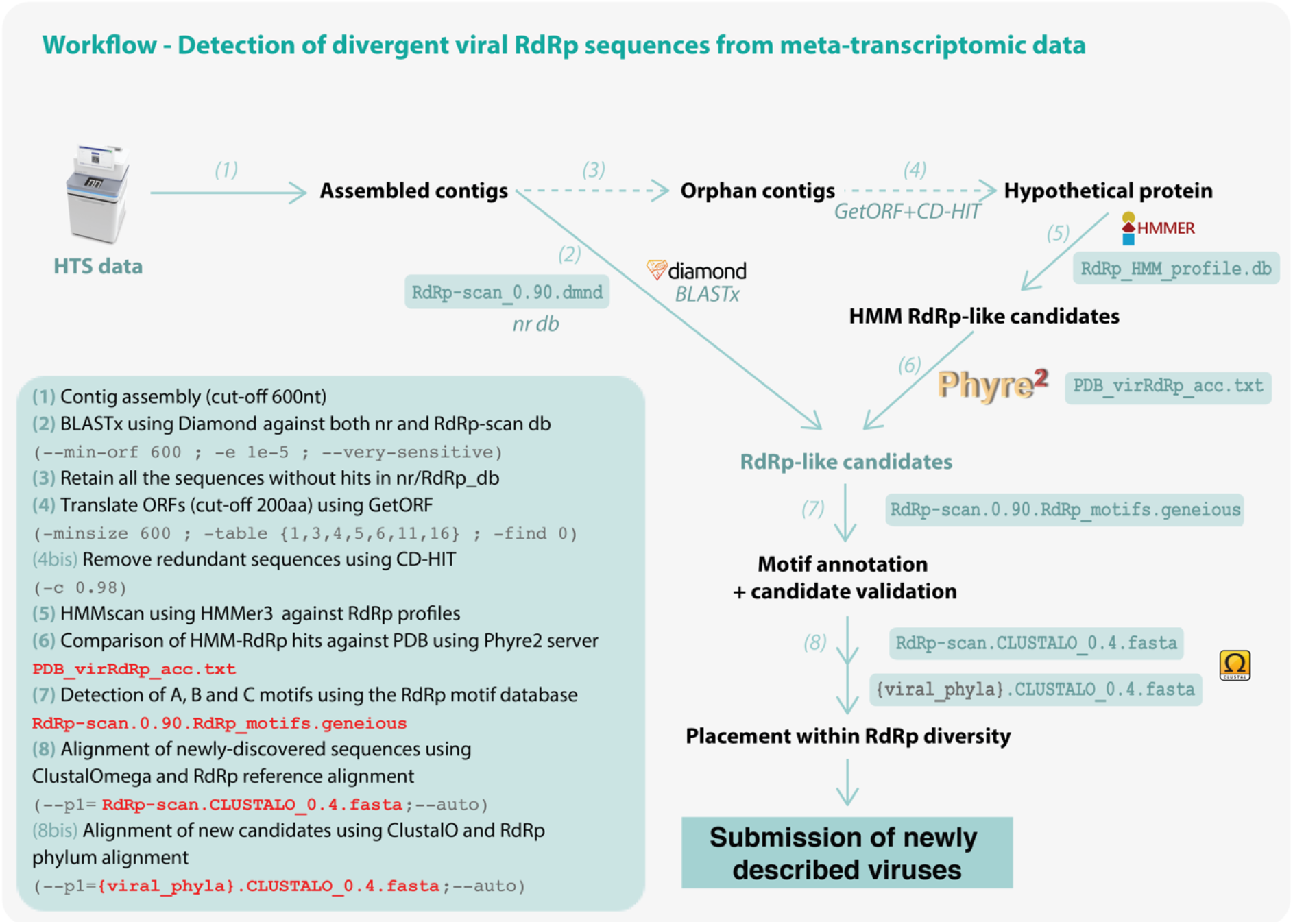
RdRp-scan workflow. **1)** Contigs are assembled from trimmed read files using assembler software using a 600nt cut-off. **2)** Assembled contigs are compared in parallel to nr and our new RdRp-scan db using Diamond BLASTx^38^, applying a *e-value* cut-off of 1e-05. All the nr matches of the sequences identified as RdRp-like are checked to remove false positive sequences, based on the e-value and BLAST scores. **3)** All “orphan” contigs - i.e., thoe without any match in nr or RdRp-scan db - are retrieved. **4)** The open reading frames or orphans contigs are then translated using getORF^46^ by applying a 200 amino acid length cut-off and according to all the genetic codes already described in viruses (i.e., translation tables 1, 3, 4, 5, 6, 11 and 16). Redundant sequences are then removed using CD-HIT ^34^ at 90% level of identity. **5)** Non-redundant candidate protein sequences are then compared to the RdRp HMM-profile database using HMMer3^41^, with an e-value cut-off of 1e-06. **6)** Retained hits are then submitted to the Phyre2 server^45^ using the batch option and results are parsed manually. To assist with this manual identification of RdRp-like hits, a list of PDB-RdRp-like structures is made available. Best hits matching with PDB non-viral entries at a confidence score higher than 90% are considered as false positives and discarded. Hits matching with RdRp PDB entries or without any confident match with PDB (at 90% confidence cut-off) are retained for further validation and characterisation steps. **7)** A, B and C motifs are then screened in the corresponding candidates using the RdRp motif database. Motif sequence and position on the sequence is manually inspected and candidate validated as true putative viral RdRp based on the presence of the three-like motifs. **8)** Confirmed candidates are then aligned with the meta-RdRp alignment using ClustalOmega^40^ and their position in the global RdRp diversity assessed using FastTREE^43^. According to their position in the corresponding tree, new RdRp-like sequences are then aligned with the closest viral phyla or order and the corresponding alignment checked and used for a more accurate phylogenetic analysis. Steps prior to the contig assembly are not mentioned.

#### Workflow to identify, annotate and assign new viral sequences

Finally, we propose a workflow that integrates the newly proposed resources as well as pre-existing open access tools, to detect new and divergent RNA virus sequences from metagenomic data (Figure 10). Briefly, the workflow consists of identifying highly divergent RdRps by combining HMMs and structural homology detection using newly built RdRp-scan RdRp and Phyre2 server, respectively. Candidates can thus be confirmed and annotated using the alignments and motif database made available in the RdRp-scan package. Importantly, the whole analysis – from the assembled contigs to the new virus RdRp annotation – can handle millions of contig sequences at a time, with relatively low computational resources.

The new viral sequences identified using RdRp-scan can be used as queries for a second round of BLASTp/HMM-profile searches which in turn may illuminate new parts of the RNA virus phylogeny. Finally, the addition of *de novo* prediction-based structures, such as those made using AlphaFold2^49^ and potentially available in AlphaFold protein structure database (https://alphafold.ebi.ac.uk/), could also be integrated to the structural-based comparison steps and are expected to enlarge the scale of structural comparison across the RNA virus phylogeny^16^.

## CONCLUSION AND PERSPECTIVES

The detection of divergent RNA viruses relies heavily on the *de novo* discovery, annotation and validation of newly described functional features such as new functional motifs and domains. By combining sequence, profile and structural-based analyses into a single workflow, our study shows that it is possible to detect RdRp sequences sharing as little as 10% sequence identity with known RdRps, and that this can be realistically conducted at a metagenomic scale. This work provides resources to ease the challenging steps that lie beyond the detection of new divergent viral sequences, particularly the identification and annotation of functional RdRp motifs and the taxonomy placement within diversity of *Riboviria*. The resources and workflow generated here will therefore facilitate the detection of divergent RNA virus sequences and expand our current knowledge of the RNA virus diversity. The recent progress in both the accessibility and accuracy of *de novo* structural predictors is also expected to provide a new perspective on the discovery of remote viral homology and will be integrated into such workflows.

## AKNOWLEDGMENTS AND FUNDING

We should like to thank Ayda Susana Ortiz-Baez and Callum Le Lay for helpful discussions and Mathilde Chagneaud for her help on designing the RdRp-scan logo. ECH is funded by an Australian Research Council Australian Laureate Fellowship (FL170100022).

## DATA AVAILABILITY STATEMENT

All the data produced in this study (alignment, database and phylogenetic tree files) are available at *https://github.com/JustineCharon/RdRp-scan*.

## SUPPLEMENTAL INFORMATION

**Figure S1.**
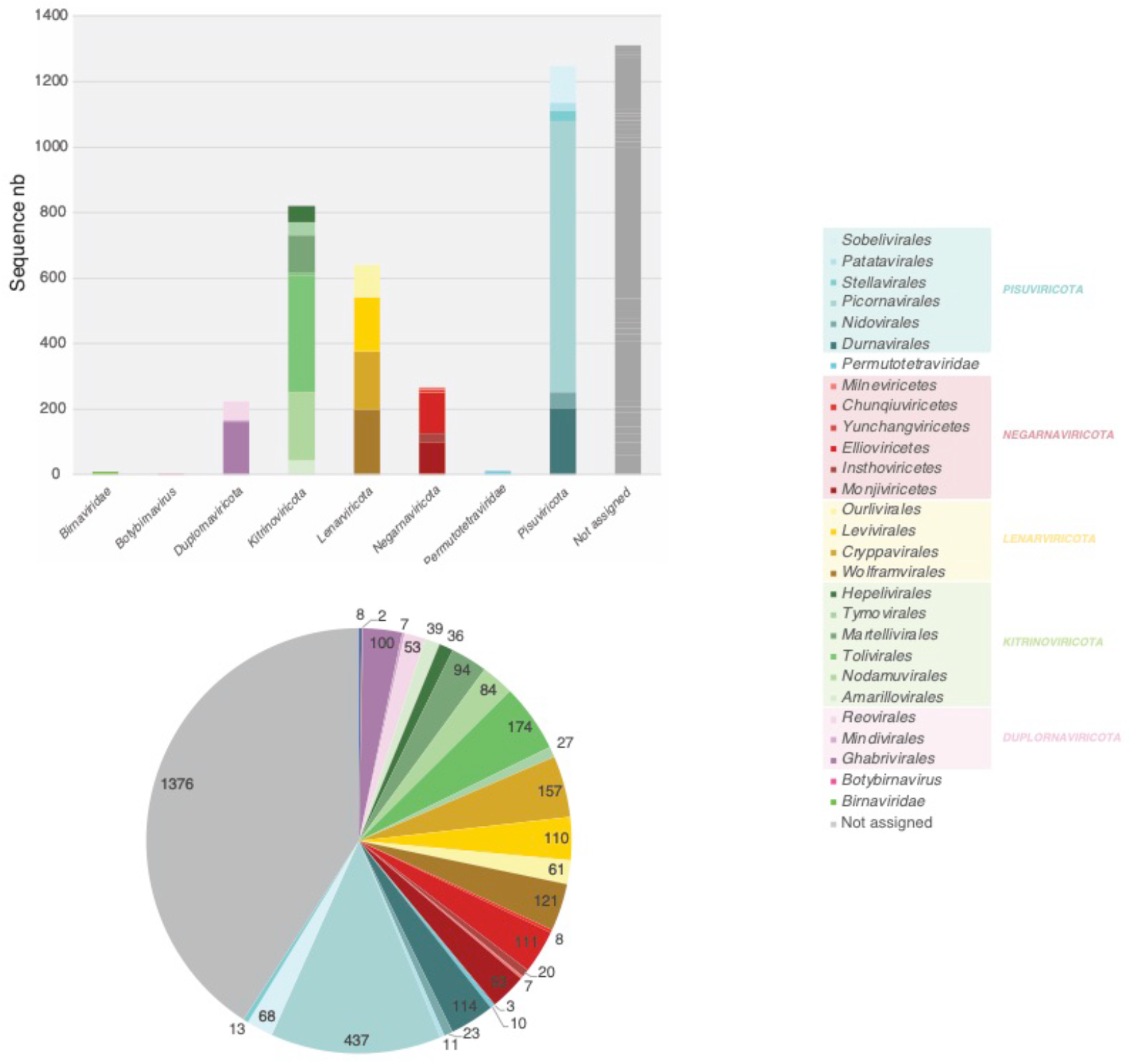
Number of sequences in the MSA used to build HMM-profile and phylogenies. Top - number of sequences in the unclassified and Riboviria phyla and order MSA used to build HMM-profiles. Bottom - number of sequences in each phyla and order MSA used to build RdRp phylogeny. Orders are coloured according to their placement into phyla (right).

**Figure S2.**
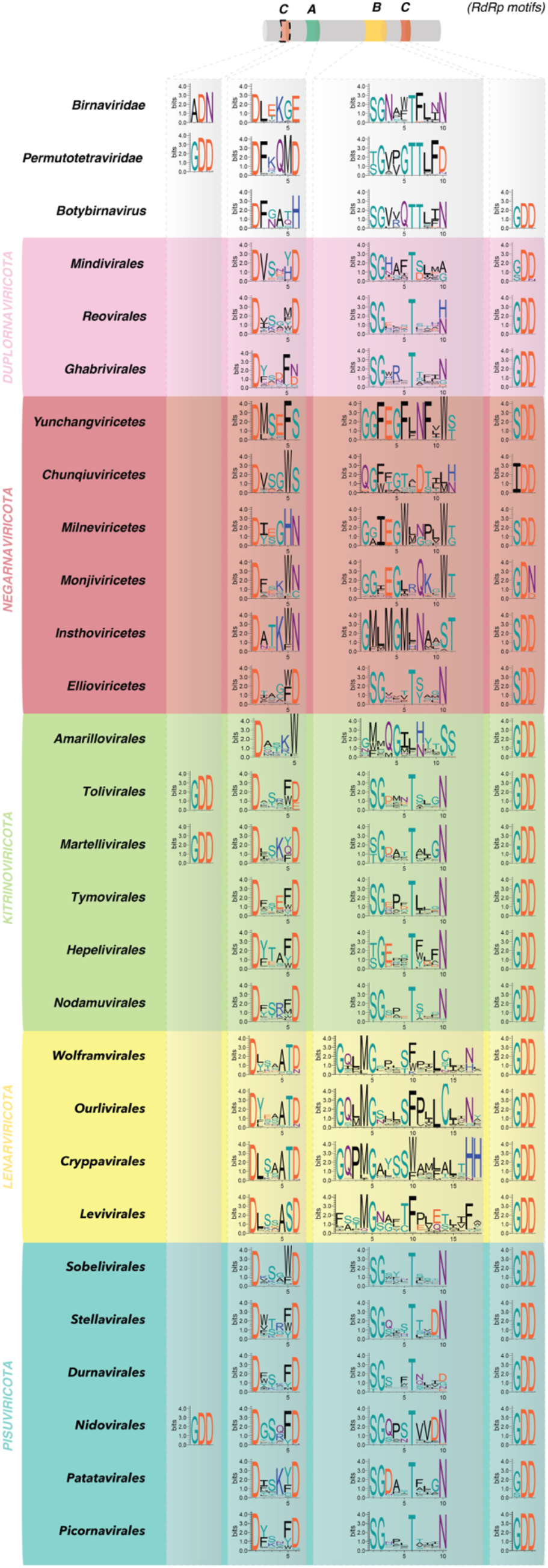
A, B and C RdRp motif diversity within the orders of *Riboviria* (*Orthornavirae*). Motifs are represented as logos obtained from alignment of RdRp within each *Riboviria* order, as defined by ICTV taxonomy and using WebLogo^42^.

